# Accumulated degeneration of transcriptional regulation contributes to disease development and detrimental clinical outcomes of Alzheimer’s disease

**DOI:** 10.1101/779249

**Authors:** Guofeng Meng, Dong Lu, Feng Yu, Jijia Sun, Chong Ding, Yan Sun, Xuan Liu, Jiapei Dai, Wenfei Jin, Weidong Zhang

**Affiliations:** Institute of interdisciplinary integrative Medicine Research, shanghai University of Traditional Chinese Medicine, shanghai, China; Wuhan Institute for Neuroscience and Neuroengineering (WINN), South-Central University for Nationalities, Wuhan 430074, China; Department of Biology, Southwest University of Science and Technology, Shenzhen, China

## Abstract

Alzheimer’s disease (AD) is extremely complex for both causal mechanism and clinical manifestation, requiring efforts to uncover its diversity and the corresponding mechanisms. Here, we applied a modelling analysis to investigate the regulation divergence among a large-scale cohort of AD patients. We found that transcription regulation tended to get degenerated in AD patients, which contributed to disease development and the detrimental clinical outcomes, mainly by disrupting protein degradation, neuroinflammation, mitochondrial and synaptic functions. To measure the accumulated effects, we came up with a new concept, regulation loss burden, which better correlated with AD related clinical manifestations and the ageing process. The epigenetic studies to multiple active regulation marks also supported a tendency of regulation loss in AD patients. Our finding can lead to a unified model as AD causal mechanism, where AD and its diversity are contributed by accumulated degeneration of transcriptional regulation.

The significance of this study is that: (1) it is the first system biology investigation to transcription regulation divergence among AD patients; (2) we observed an accumulated degeneration of transcription regulation, which well correlates with detrimental clinical outcomes; (3) transcriptional degeneration also contributes to the ageing process, where its correlation with ages is up to 0.78.

## 1 Introduction

Alzheimer’s disease (AD) is a complex chronic neurodegenerative disease that has been intensively studied for decades. However, its causal mechanismd remain elusive [1]. More attentions have been focused on the visible neuropathological features, especially the amyloid plaques and neurofibrillary tangles. Amyloid plaques are composed of depositions of insoluble and densely packed amyloid beta (A*β*) protein whereas neurofibrillary tangles are composed of aggregations of hyperphosphorylated tau protein [2]. Early-onset AD studies in rare families led to the discovery of three genes, amyloid precursor protein, presenilin 1, and presenilin 2 that demonstrated the causal effects of A*β* in the AD progression [3]. However, this is challenged by the observation that some patients with substantial accumulations of plaques have no cognitive impairment [4] and that drugs targeting A*β* all failed in clinical studies [5]. Although neurofibrillary tangles have a stronger correlation with the decline of cognitive ability and have drawn more attentions recently [6, 7], strong or direct evidence linking neurofibrillary tangles to AD is still lacking [8].

Integrated systematic approaches, especially coexpression regulatory network analysis, have advanced our understanding of AD in different ways [9, 10, 11]. In such studies, biological networks are constructed using gene-expression data to identify the gene modules related to AD genesis and development by integrating quantitative evaluation into both AD neuropathology and cognitive ability decline. In our previous work, we studied transcriptional dysregulation between AD patients and control subjects, and identified a core network [12]. Combining computational prediction with experimental perturbation allows the discovery of causal pathways and regulators, which can be used for therapeutic targets. Compared to studies performed at the single gene level, network-based analysis provides a more comprehensive insight into AD.

The commonly used algorithms for network analysis, such as WGCNA [13], MEGENA [14] and SpeakEasy [15], apply coexpression approaches to identify gene modules with common regulations or biological involvements. Originally, network analysis was often limited by the cohort size and/or less realization to the complexity of diseases. Therefore, the existing tools usually take less consideration to patients’ diversity and assume all the subjects under the same regulation patterns. Recent edivdences suggest that AD patients have great diversity at both neuropathologic burden and disease development, mainly contributed by the divergent involvement of AD genesis mechanisms [16]. With the advancement of the scientific community for AD studies (e.g. the AMP-AD project), large cohorts have been collected for integrated analysis. Such advancements persuade us to rethink about the complex dysregulation mechanisms among the AD patients.

Recently, epigenetic studies of AD patients have enhanced our understanding of the dysregulation occuring during AD genesis [17, 18, 19, 20]. Investigation of DNA methylation marks and maintenance factors reported decrements of DNA methylation in AD patients [21]. Genome-wide studies of CpG islands identified altered DNA methylation and suggested their impacts on the AD risk genes [22]. Histone modification studies to active epigenetic marks, such as H4K16ac, H3K9ac and H3K27ac, suggested that abnormal epigenetic regulation affects the regulation of AD genes [23, 24]. Additionally, recent studies reported that large-scale changes in H3K27ac could be driven by tau pathology in human brains [19] and that HDAC3 inhibition could reverse AD-related pathologies in the animal model of AD [25]. These finding suggests that there is a close cross-talk between gene regulation and AD genesis. However, there are still gaps in our understanding between abnormal epigenetic regulation and the causal mechanisms of AD. Many efforts were put on the known AD genes, such as APP, MAPT and GSK3B [26]. Similar to genome-wide association studies, it is easy to identify factors associated with AD genesis but difficult to elucidate the complex causal mechanisms in an integrated framework or to predict disease clinical outcomes of AD patients.

In this work, we performed a two-stage study to reveal the divergence of transcriptional regulation among AD patients. In the first stage, we utilized a computational method to study transcription factor (TF)-mediated regulation in a large cohort of subjects, including both AD patients and normal individuals at different clinical stages. We found that transcriptional regulation tended to get weakened or missed in AD patients, and some of these regulation loss were closely associated with clinical features. Interestingly, regulation loss almost indicated detrimental clinical outcomes. Functional annotations further confirmed that regulation loss disrupted the AD-related biological processes, such as protein degradation, neuroinflammation, mitochondrial dysfunction, and neuronal/synaptic function. To measure its effects, we came up with a new measurement, regulation loss burden (RLB), to describe the accumulated degree of regulation loss and found that RLB better indicated detrimental clinical outcomes than the existing methods. In the second stage, we performed genome-wide studies to active epigenetic marks, including histone modification marks, open chromatin accessibility and three TF binding sites. A strong tendency of active mark loss was observed in AD patients, which was consistent with our computational results. Overall, our results suggest critical roles of accumulated transcriptional regulation loss in AD development and clinical outcomes. It could lead to a unified model to elaborate the complex causal mechanisms of AD and the diversity among AD patients (see Figure 1).

**Figure 1:**
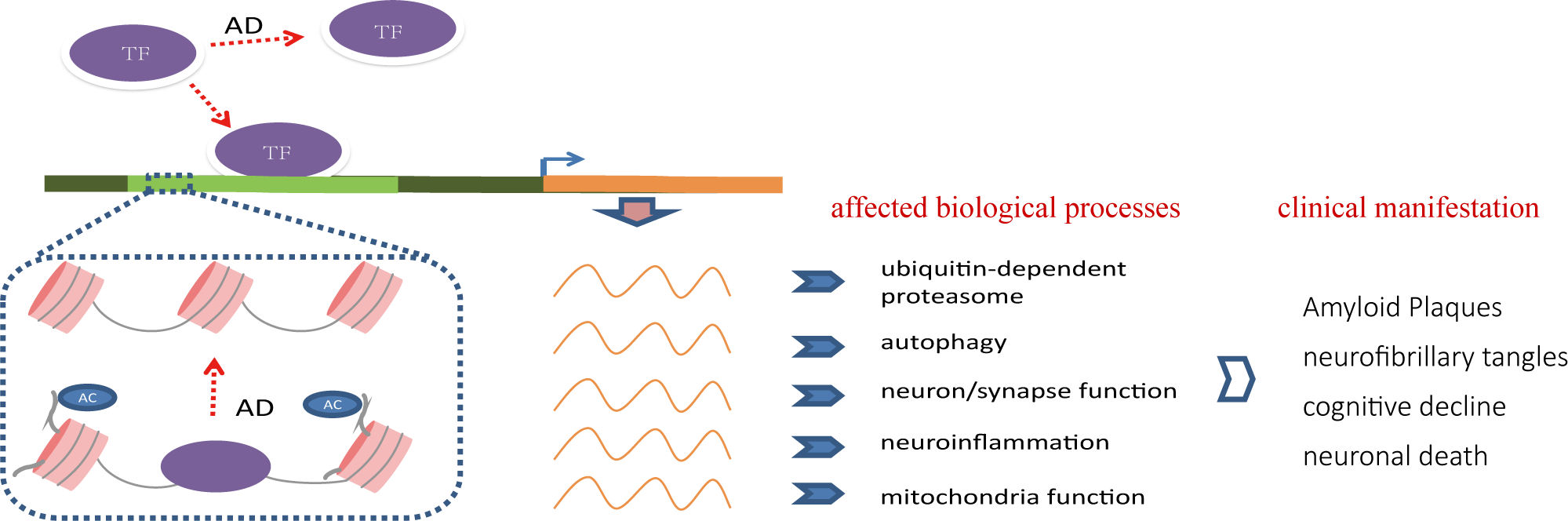
Accumulated degeneration of transcription regulation contributes to AD development and detrimental clinical outcomes. Our results suggest that transcriptional regulation tends to get lost in AD patients, which disrupts the normal cellular function of brain, e.g. protein degradation, neuroiflammation, mitochondrial, neuronal/synaptic function, and contributes to the detrimental clinical outcomes. This finding may lead to a unified model to elaborate the causal mechanisms of AD, where brain transcriptional regulation degenerates from an organized system in normal individuals into a deficient system in the AD patients.

## 2 Results

### 2.1 Weakened and missed regulation widely exists in AD patients

We explored the divergence of transcriptional regulation among AD patients. Therefore, we developed a bi-clustering algorithm to study TF-mediated regulation in patients at different clinical stages. The design of this algorithm is based on two assumptions: (1) that we can find a set of biomarker genes to indicate the TF regulatory activity; and (2) that AD patients can be clustered into groups with different TF regulation status. As shown in Figure 2(a,b), this algorithm takes only expression data as the input and then output a subset of patients that is regulated by a specific TF. Active TF regulation is identified if it satisfies following three criteria: (1) TF-gene co-expression correlation |*r*| is greater than 0.8, which is a strict cutoff to identify biomarker genes indicating TF regulatory activity; (2) the selected patient subset has more than 50 patients; (3) TF strictly regulates least 30 genes. In case if no TF satisfies the cutoff of |*r*| > 0.8 in any subset of patients, the TFs will be assigned with a type of “non-dominant regulation” (NR). NR regulators are not considered to have no clear regulatory role. The regulatory types of other TFs are determined by their regulation strengths in the remaining patients. For example, “DR” is assigned if the |*r*| of remaining patients satisfies *r* > 0.6, which indicates that such a TF has a dominant regulatory role in all patients. “WR” and “MR” indicate a weakened or missed regulation in the remaining patients when |*r*| is above or below 0.3. The patients with weakened or missed regulation are supposed to have regulation loss.

**Figure 2:**
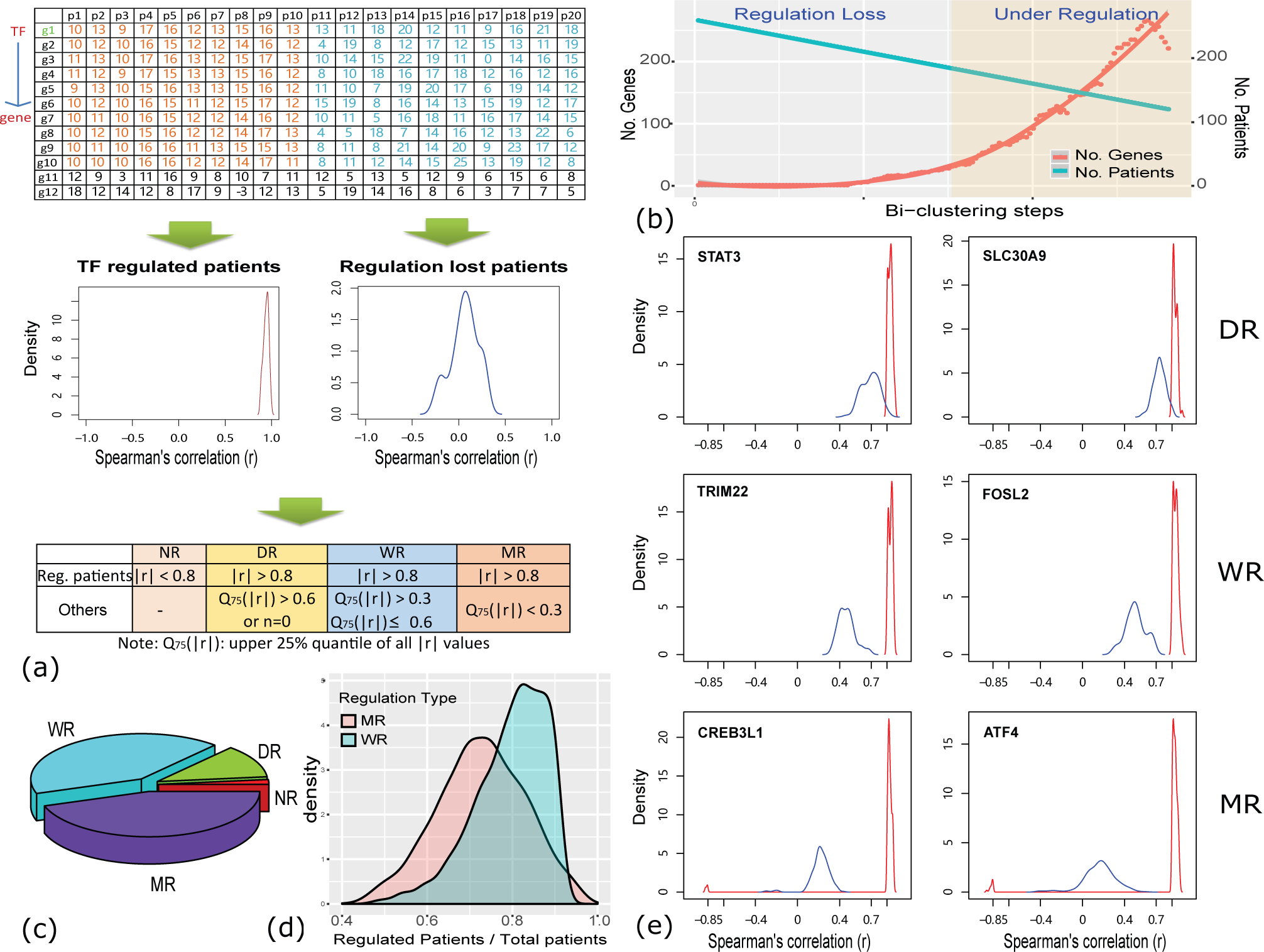
Transcription factor-mediated regulation loss widely exists in AD patients. (a) A computational method to discover the missed and weakened regulation. In this process, a bi-clustering algorithm is applied to cluster the patients into two groups with or without TF regulation. Based on the strength of regulation loss, TFs are assigned with types of no dominant regulation (NR), dominant regulation (DR), weakened regulation (WR) or missed regulation(MR); (b) the dynamic curve of gene and subject number during the bi-clustering analysis, reflecting the regulatory activity of studied TFs; (c) the distribution of predicted regulation types, where MR and WR are most observed; (d) TFs tends to have regulation loss in only portion of subjects (about 25-40% of studied subjects); (e) TF-gene correlation distribution of 6 exemplary regulators, where TFs take dominantly regulatory roles in some subjects (red line) while their regulations are weakened or missed in other patients (blue line), indicating existence of regulation loss among the patients.

Before application, we performed four evaluations on its reliability, including the following (1) the ability to identify the subset of patients with different TF regulations; (2) false positive ratio of bi-clustering prediction; (3) the impacts of different correlation cutoffs on the analysis results; and (4) evaluation using independent normal brain tissues. Our evaluations suggested that the predicted regulation loss was not due to technical biases of strict cutoffs and that bi-clustering analysis could recover the true regulation loss of AD patients (see details in Supplementary Results). Meanwhile, we also found that |*r*| > 0.8 was a reasonable cutoff to achieve a good analysis power for the data used in this work.

We analyzed the RNA-seq expression data for 945 autopsied samples from 364 subjects in four brain regions: frontal pole (BA10), superior temporal gyrus (BA22), parahippocampal gyrus (BA36), and frontal cortex (BA44). These subjects had diverse clinical manifestations, e.g. cognitive score and braak stages. Among them, approximately 61% were diagnosed as having pathological AD or probable AD (see Figure S1 for detailed clinical information) [27]. Specifically, 869 brain-expressed TFs were selected to study their regulatory status among these subjects (see Methods for detail). Bi-clustering analysis identified only nine TFs, including STAT3, ST18, CSRNP3, LMO4, CNOT7, SLC30A9, PEG3, SUB1 and MEF2C, taking strict regulation in all the subjects at a cutoff of |*r*| > 0.8. To reduce the false negative discovery of DR TFs due to an excessively strict cutoff, we gradually loosed the correlation cutoff to the 75% quartile of |*r*|, *Q*_75%_(|*r*|) = 0.6 or the minimum number of co-regulated genes to 5. More regulators were identified and they were assigned with regulatory types of “DR”. For other TFs, the bi-clustering algorithm was optimized to select the maximum number of genes that satisfied a cutoff of *r* > 0.8. By calculating the coexpression correlation in the remaining subjects, we found that approximately 40% of TF had weakened regulatory roles in the remaining subjects (0.3 < *Q*_75%_(|*r*|) < 0.6) (see Figure 2(c)). Therefore, these TFs were assigned with the regulatory types of “WR”. Meanwhile, another 40% of TFs were assigned with the regulatory type of “MR”, where *Q*_75%_(|*r*|) was less than 0.3 in the remaining subjects. As shown in Figure 2(d), we found that more than half of the subjects were under the TF regulation when the minimum TF-regulated genes were greater than 30. We also checked the regulatory relationship between predicted TFs and subjects. We found that the regulation loss was not specific to any subset of subjects but widely existed in all subjects. Meanwhile, any subject could be under missed or weakened regulation of multiple TFs (See Figure S2). Figure 2(e) showed the regulatory status of some exemplary TFs, where subjects were clustered into two subsets under different TF regulations, e.g. DR, WR and MR. We also performed TF over-representation analysis to check if the TF target genes were enriched with TF-binding motifs. Using the annotation of RcisTarget [28], we selected 487 TFs for evaluation and found that 31% of them were enriched with corresponding TF binding motifs (see Table S1), which was comparable to our previous findings [29, 30]. This result suggested that TF-gene regulation identified by bi-clustering analysis was more likely be bound by predicted TFs.

We speculated that the decreased gene expression of TF genes contributed to the regulation loss. Hence, we evaluated the differential expression statuses of WR and MR regulators and found that only about 10%-15% of TF genes displayed significant expression differences between subjects with or without regulation loss at a cutoff of *p* < 0.01 (see Table S2), suggesting that most of regulation loss was not related to decreased expression of TF genes. We further explored the transcript isoform usage of undifferentially expressed regulators using RNA-seq data [31]. However, we failed to find a strong switch in isoform usage, especially for the abundantly expressed isoforms. Overall, it seems that regulation loss is beyond expression changes of regulator genes. We also investigated the impacts of neuronal loss based on brain-specific marker genes and did not find any evidence for the existence of neuronal loss (see Supplementary Results).

### 2.2 Regulation loss almost indicates detrimental clinical outcomes

Next, we investigated if the regulation loss was associated with AD related clinical features. Based on predicted regulatory statuses, the subjects were automatically clustered into two non-overlapping groups: the subjects under TF regulation and the ones with regulation loss. Three clinical traits, including the cognitive score (CDR), Braak score (braak) and amyloid plaque mean size (plaque) were checked for clinical feature differences between the two groups using Kolmogorov–Smirnov (K-S) tests. To control for false prediction, we estimated the false discovery ratio (FDR) by randomly shuffling clinical trait values. At a cutoff of *p* < 0.01 and FDR < 0.05, 291, 277, 335 and 159 TFs were predicted to have association with at least one of three clinical traits, accounting for about 38%, 36%, 47%, and 22% of MR/WR TFs in BA10, BA22, BA36 and BA44 regions, respectively (see Figure 3(a) and Table S3). To further evaluate the validaity of these observations, we performed another round of simulation evaluation by repeating the same analysis using random sample combinations. Under this setting, the maximum number of clinical associated TFs was less than 10 for all three clinical traits and the possibility for our observation was nearly impossible (*p* = 0), which suggested a good confidence to trust the association between regulation loss and clinical traits. Even though three clinical traits were studied, we found that there was always more association with CDR and plaque scores than braak (see Figure 3(a)), which is consistent with published reports [10]. We also noticed that WR regulators were more associated with clinical outcomes than MR regulators (see Figure S3).

**Figure 3:**
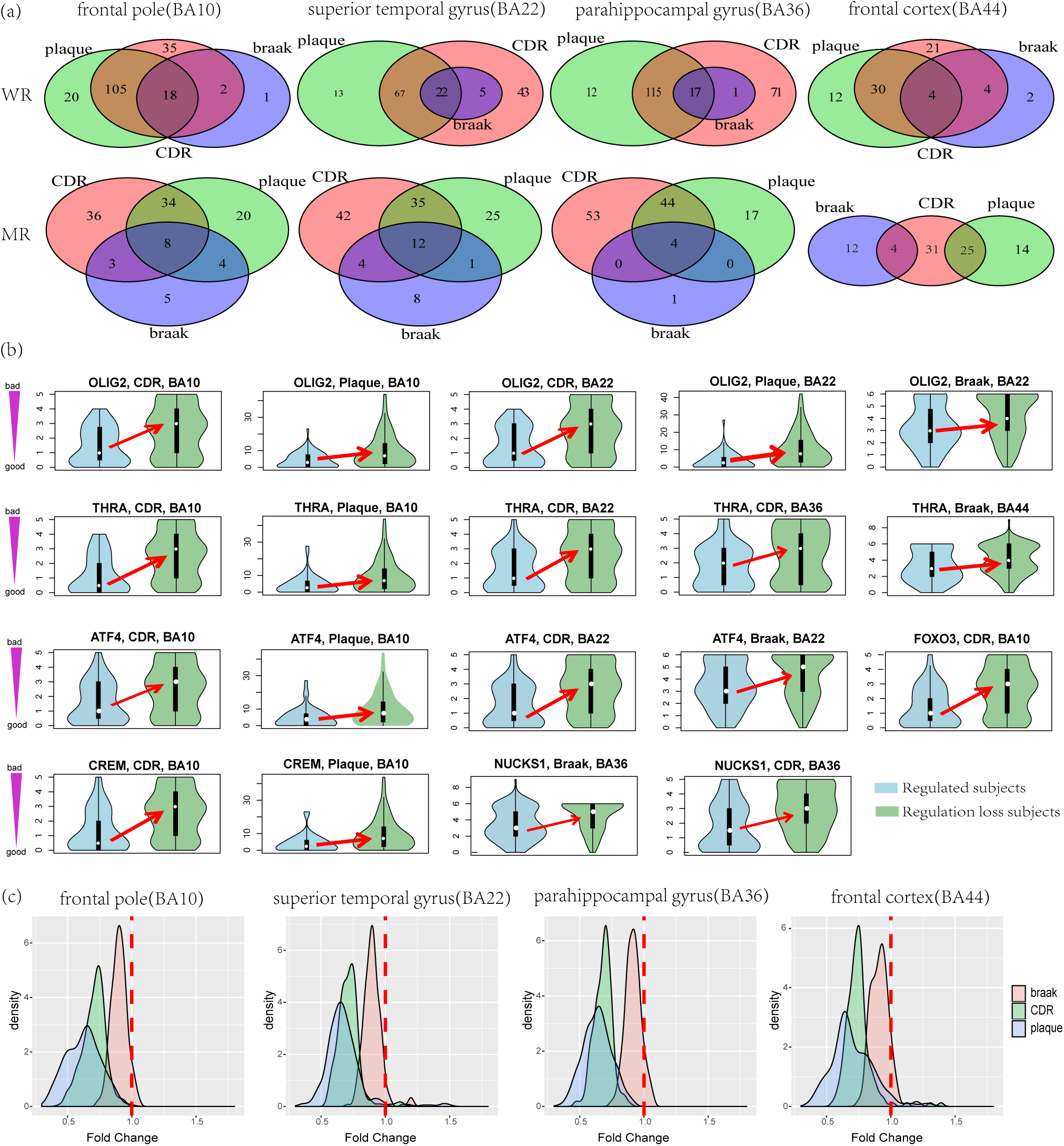
Regulation loss almost indicates detrimental clinical outcomes. (a) WR and MR TFs predicted with clinical association in four brain regions. Three clinical traits are used, including cognitive score (CDR), braak score (braak) and amyloid plaque mean size (plaque). Four brain regions are studied: frontal pole (BA10), superior temporal gyrus (BA22), parahippocampal gyrus (BA36) and frontal cortex (BA44). (b) The clinical association of exemplary TFs. We observed that subjects with regulation loss usually had worse clinical outcomes. (c) Regulation loss is always associated with detrimental clinical outcomes. In this study, we identified about 250 TFs in each brain region to have association with at least one AD-related clinical trait; fold change analysis indicated that regulation loss of those TFs almost indicated detrimental clinical outcomes. Here, lower fold changes (< 1) indicate changes to the more detrimental clinical outcomes.

Figure 3(b) shows representative TFs and their clinical associations. Many of them have been reported for their involvement in AD or brain function. For example, OLIG2 (Oligodendrocyte transcription factor) locating on chromosome 21 in a region contributing to the cognitive defects of Down syndrome [32]. Evidences suggest that OLIG2 participates in fate switch of neurons in AD patients [33]. In our result, OLIG2 was an MR regulator and was significantly associated with both CDR (*p* = 6.86*e* − 7) and Plague (*p* = 7.19*e* − 9) in three brain regions, including BA10, BA22 and BA44. THRA (Thyroid Hormone Receptor, Alpha) is a MR regulator associated with CDR (*p* = 2.02*e* − 8) and Plague (*p* = 1.04*e* − 6) in two brain regions. It is essential for normal neural development and regeneration [34]. THRA has been reported to have a weak genetic link with AD [35]. ATF4 has been widely reported for the increased expression in AD patients and transcriptional mediator roles in neuron degeneration, metabolism and memory formation [36]. FOXO3 has been reported in genetics meta-analysis [37] and participating in neuronal mitophagy [38] and insulin-like growth factor I signaling pathway [39]. CREM has been reported to have increased gene and protein levels in AD [40]. NUCKS1 is a risk gene for Parkinson’s disease [41].

Using these exemplary TFs (see Figure 3(b)), we observed consistent clinical associations, where TF regulation loss was observed more often in the patients with a worse diagnosis. We further extended our analysis to other AD associated TFs by checking the fold change of clinical traits. As shown in Figure 3(c), nearly all tested TFs tended to be associated with detrimental clinical outcomes, such as declined cognitive ability, increased Braak stage and increased amyloid plaque size. The similar tendency was consistently observed in four brain regions. Among the three clinical traits, CDR and plague always indicated more clinical association, e.g. more associated TFs or larger fold changes. Considering that the bi-cluster analysis was blind to clinical annotations, it seems that TF regulation loss, except for very few TFs, always had a consistent impact on the clinical outcomes of patients. We also investigated the few TFs with inconsistent clinical association and found that they usually displayed less significance for clinical association (see Table S3). Overall, these findings suggests that regulation loss is closely associated with detrimental clinical outcomes of AD patients.

### 2.3 Transcriptional regulation loss disturbs the AD-related processes

Our algorithm predicted 6300 biomarker genes to be regulated by at least one WR or MR TF in any of four brain regions. Among them, 43% of genes were regulated by only one TF and 80% were regulated by less than 5 TFs (see Figure S4). It seems that most of the selected genes were dominantly regulated by only one or a few TFs, which is consistent with the assumption for biomarker genes. Therefore, it is feasible to study the functional involvement of TFs by enrichment analysis of their regulated genes. Figure 4(a,b) highlights the key pathways and biological processes associated with regulation loss. Even though the four brain regions were investigated independently, they indicated similar functions.

**Figure 4:**
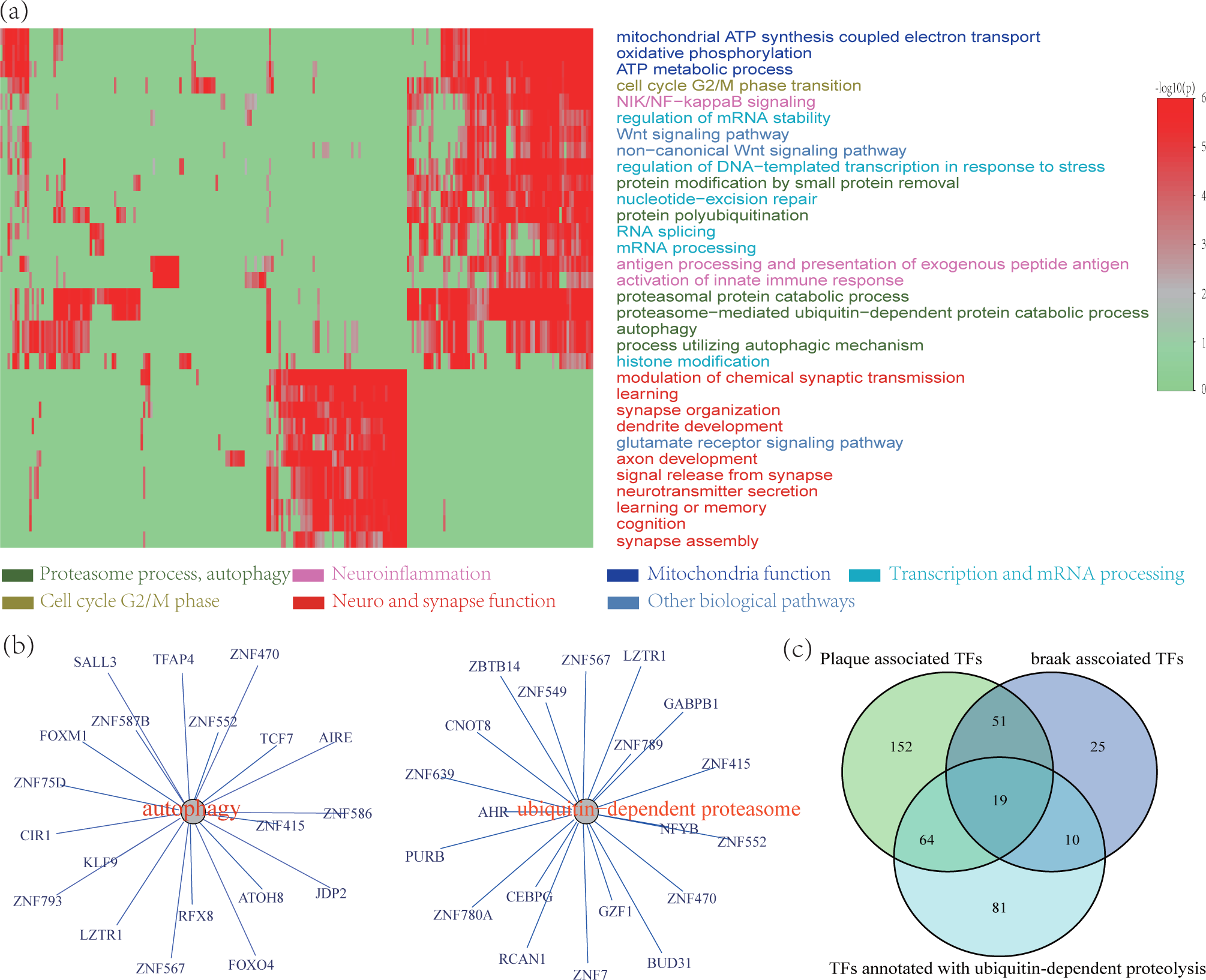
Functional involvement of regulation loss. (a) The functional involvement of regulation lost TFs was annotated by functional enrichment analysis to their downstream genes. The similar results were observed in four brain regions, such as protein degradation, neuroinflammation, mitochondrial function, neuronal or synaptic functions. (b) The top 20 TFs associated with protein degradation process and synapse function. (c) TFs annotated with ubiquitin-dependent proteolysis are more associated with plague and braak scores (*p* < 0.005).

Among them, protein degradation, especially the ubiquitin-proteasome system (UPS), was the most affected biological process (see Figure S5). Its enrichment was observed in more than 50% of TFs in all four brain regions. The abnormal accumulation and misfolding of A*β* and tau are two features of AD patients as the extracellular amyloid plaques and intraneuronal neurofibrillary tangles [42]. Growing evidences support a tight link between the impairment of the UPS and it is speculated that the age-dependent impairment of UPS the could affect the degradation of A*β*, which leads to abnormal accumulation and aggregation in the brain; A*β* can also inhibit the UPS activity [43]. We found that more than half of the TFs annotated with ubiquitin-dependent proteasome function were also associated with plaque and braak scores (*p* < 0.005) (see Figure 4(c)), suggesting that amyloid plaque and neurofibrillary tangles are closely associated with dysregulated protein degradation. Additionally, autophagy, another protein degradation system [44], was enriched in the downstream genes of more than 60 TFs, among which some have been reported for their regulatory roles in autophagy, such as CEBPG, E2F1, HLTF, HSF2, KDM1A, NFE2L1, NR1D1 and TRIM28.

Neuroinflammation was the second most enriched biological process. Terms, such as “NIK/NF-kappaB signaling”, “innate immune response activating cell surface receptor signaling pathway” and “antigen processing and presentation of exogenous antigen” are enriched with about 40% of TFs in all four brain regions. This finding is consistent with the reports that activation of inflammatory system contributes to AD pathogenesis [45, 46]. Mitochondria related function, such as “electron transport chain” and “oxidative phosphorylation”, were enriched in about 40% of TFs. Extensive literature and evidence support the role of mitochondrial dysfunction and oxidative damage in the genesis of AD [47]. Additionally, mitochondria dysfunction is also related to other AD-related processes. For example, age-dependent oxidative stress induces the accumulation of A*β* and the deposition of neurofibrillary tangles [48]. The cell cycle G2/M phase transition was enriched in about 40% of TFs. Neurons suffering from synaptic dysfunction, oxidative stress or other stress factors may enter cell cycle and this is linked to tau hyper-phosphorylation and A*β* [49]. Most the reports on the cell cycle are related to the G0 phase and it is still not clear why only G2/M phase transition terms are enriched in our analysis. Neuronal and synaptic function related processes were enriched with about 20% of TFs. Neuronal and synaptic losses occur in the entrie phase of AD and they are correlated with severity cognitive declines [50]. Synaptic function is closely associated with proteasomal processes. For example, the ubiquitination of synaptic receptors and kinases has crucial roles in synaptic transmission and plasticity [51]. Oligomeric A*β* has toxic effects on synaptic function. Other enriched processes include Wnt signalling pathway, rRNA processing, histone modification and other processes.

Overall, our functional studies of WR/MR TFs successfully re-discovers the key processes and pathways related to AD genesis and development. Correspondingly, these findings suggest that regulation loss is involved in AD genesis and development by affecting AD-related processes, especially protein degradation.

### 2.4 Regulation loss burden better correlates with AD clinical outcomes

We next investigated if regulation loss could be used as a molecular marker to predict clinical outcomes of AD patients. Thus, we introduce a new measurement, regulation loss burden (RLB), which describes the accumulated degree of transcriptional regulation loss in AD patients (see Methods section). Unlike other measurements, it reflects overall transcription regulatory activities in one patient.

In the first analysis, we used all the WR, MR and DR TFs as input to calculate the RLB and found that RLB had strong correlations with all three clinical traits, especially CDR and Plague (see Figure S6). Spearman’s correlations with CDR were calculated to 0.39 in BA10 and BA36. Other clinical traits also reached significant correlations (*p* < 0.01). Considering that many TFs are not related to AD, we further restricted TFs to the these associated with AD. We observed that the association was greatly improved for all three clinical traits (see Figure 5(a)). In BA10 and BA36, the RLB had the best correlation with CDR, where Spearman’s correlation was *r* = 0.5. In BA10, RLB had the best correlation with braak score at *r* = 0.48. In other brain regions, RLB also achieved good correlations with CDR and Plaque, too. As an evaluation, we also investigated the relevance with other covariates, such as age and sex. As shown in Figure 5(b), RLB only exhibited a strong correlation with AD-related clinical traits.

**Figure 5:**
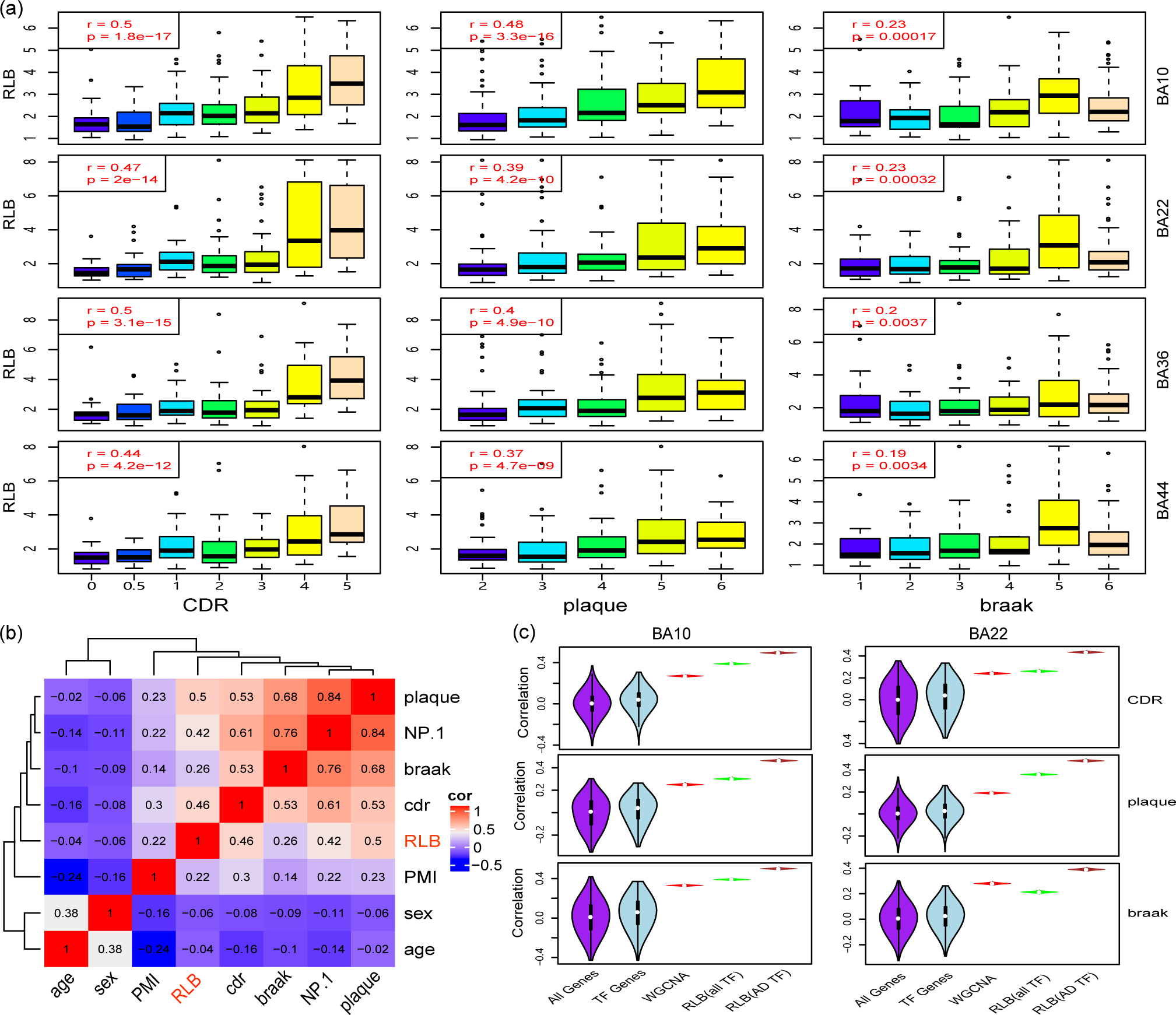
Regulation loss burden better correlates AD clinical outcomes than existing methods. (a) Association between regulation loss burden and clinical outcomes in four brain regions, where the correlation value is up to *r* = 0.5; (b) Regulation loss burden is only associated with AD-related clinical traits. PMI: Post-Mortem Interval; NP.1: Neuropathology Category. (c) Regulation loss burden better indicates the AD development than genes, transcription factors and WGCNA modules (see Figure S7 for more results). Here three clinical traits, including CDR, Plaque, Braak scores, are used.

Transcriptomic features have been previously explored for their associations with clinical features [11]. However, most of these studies failed to identify genes or modules with strong correlations to AD clinical traits. We compared RLB with these measurements (see Figure 5(c) and Figure S7). At the single gene level, no gene displayed a better association than RLB, e.g. *r* > 0.5. The maximum correlation was observed with Plaque score for HMBOX1 at *r* = 0.44. For CDR and Braak score, the maximum correlation was with ETV1 at *r* = 0.43 and with VGF at *r* = 0.40, respectively. Among the TF genes, except ETV1, no other TF gene had a correlation value greater than 0.4. Next, we performed WGCNA network analysis in each brain region and the predicted modules were evaluated for all clinical traits. Without including the grey module, the maximum correlation value (*r* = 0.34) was observed between CDR and a module predicted in the BA36 region, which was lower than that of RLB. Overall, our evaluation suggests that RLB better correlates with AD clinical outcomes than the measurement using single gene or gene modules.

### 2.5 Regulation loss in independent datasets

We repeated the same analysis using five other set of expression data collected from the published studies. They included ROSMAP expression data using microarray [9] and RNA-seq [10], HBTRC microarray study [9], Mayo’s RNAseq study for cerebellum (CBE) and temporal cortex (TCX) [52]. The dataset from Mayo and HBTRC had no extra clinical annotations and only a binary disease status was available. Thus, the RLB values were checked between the AD patients and control subjects. As shown in Figure 6(a-c), AD patients displayed more regulation loss burden than control subjects. Among them, the HBTRC dataset had more subjects (463 subjects), which allowed a more reliable prediction of regulation loss. Specifically, 196 TFs were identified to be associated with AD disease status, and statistical test suggested a significant regulation loss in AD patients (*p* = 3.8*e* − 54). The ROSMAP dataset included both microarray and RNA-seq data. The analysis results using the microarray data indicated a positive correlation between an increased RLB value and detrimental clinical outcome (see Figure 6(d, e)). However, using the ROSMAP RNA-seq data, the RLB values did not indicate a consistent tendency. As shown in Figure 6(f), a positive correlation was observed when Braak score was less 3 and a negatively correlated tendency was observed when Braak was greater than 3. It seems that the ROSMAP RNA-seq dataset only supports the hypothesized relationship between RLB and clinical outcomes at an early stage of AD. We evaluated the ROSMAP RNA-seq data but did not find any clue for inconsistency. Considering the analysis results using microarray, we still believe regulation loss to be associated with AD clinical outcomes.

**Figure 6:**
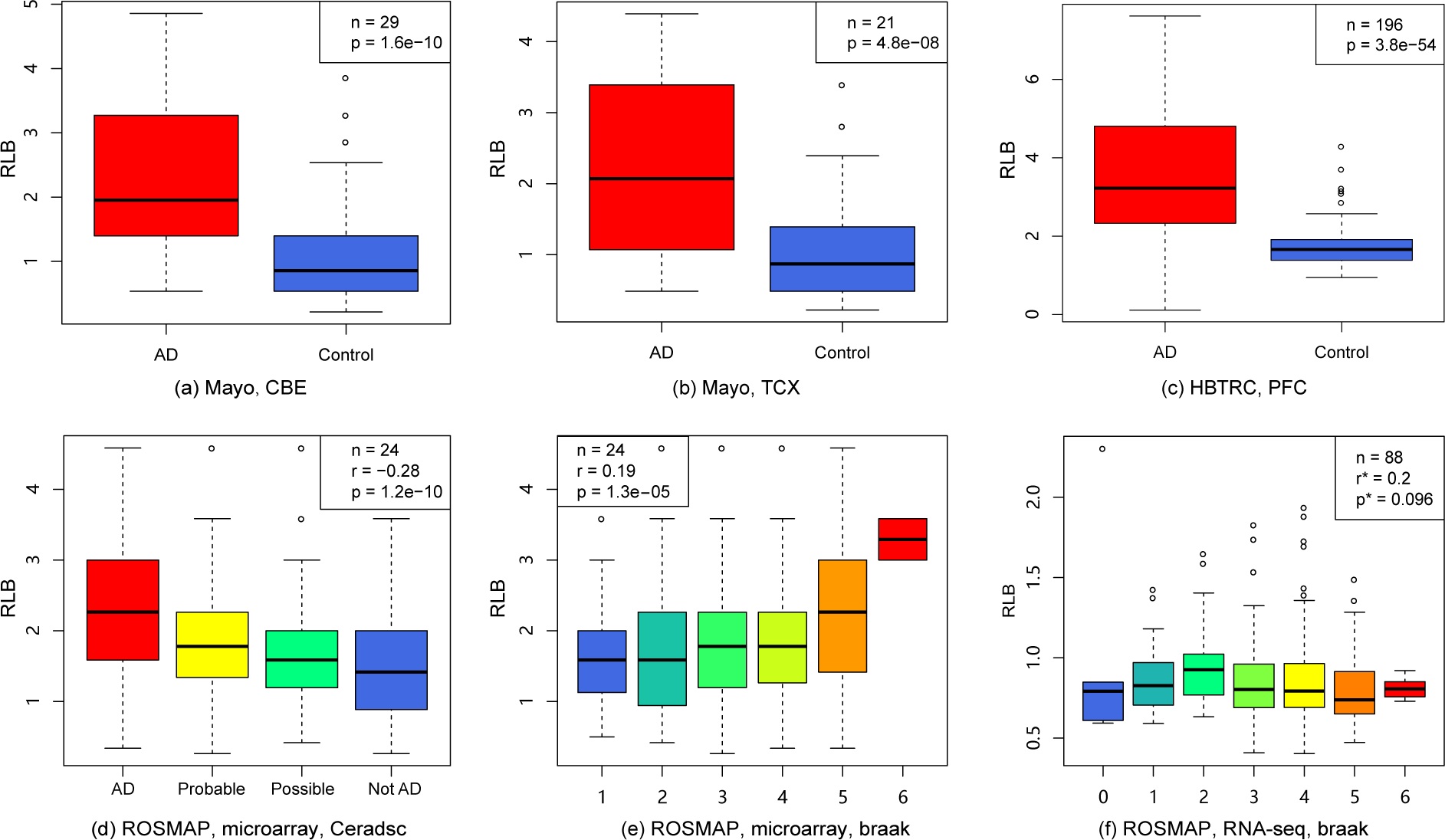
Regulation loss is associated with the detrimental AD clinical outcomes in five independent datasets. (a)Mayo’s data for cerebellum; (b)Mayo’s data for temporal cortex; (c) HBTRC study; (d) ROSMAP microarray study, association with CERAD score; (e) ROSMAP microarray study, association with braak; (f) ROSMAP RNA-seq study, association with braak (*r* and *p* is calculated using only braak=0,1,2).

Overall, five independent datasets partially or completely support the same conclusion that regulation loss correlates with the detrimental clinical outcomes of AD patients.

### 2.6 Loss of active epigenetic regulation marks in AD patients

We evaluated the transcriptional regulation status of AD patients by genome-wide screening to the active regulation marks, including histone modification, open chromatin accessible regions, transcription factor binding sites. In this study, we used two strategies to study their statuses. The first option is to do differential peaking analysis to study the peak intensity between AD and normal samples. Another one is to count absolute number of peaks among the AD patients and normal samples.

H3K27ac is an active regulation mark and is locatd at both proximal and distal regions of transcription start site (TSS) [53]. H3K27ac has been observed to display differential enrichment in AD patients, especially in the regulatory regions of AD risk genes [24]. We performed peak calling analysis on H3K27ac ChIP-seq data and found fewer peaks in AD patients compared to that of controls, suggesting an active regulation loss in AD patients (see Figure 7(a)). Applying a modified analysis pipeline origninally introduced in [24], we re-evaluated H3K27ac enriched regions and observed a strong tendency of H3K27ac mark loss in AD patients, where the ratio of loss against gain was 1:0.37 (see Figure 7(b)). Then, we checked if the H3K27ac loss was associated with dysregulated gene expression. Using the downstream target genes of the top 10 dysregulated TFs (see Table S3), a significant overlap was observed between dysregulated genes and the genes with H3K27ac loss in the promoter regions (*p* = 5.08*e* − 32 by Fisher’s exact test) (see Figure 7(c). Similar but weak results were observed with H4K16ac, which is another active regulation mark [23] (see Figure S8).

**Figure 7:**
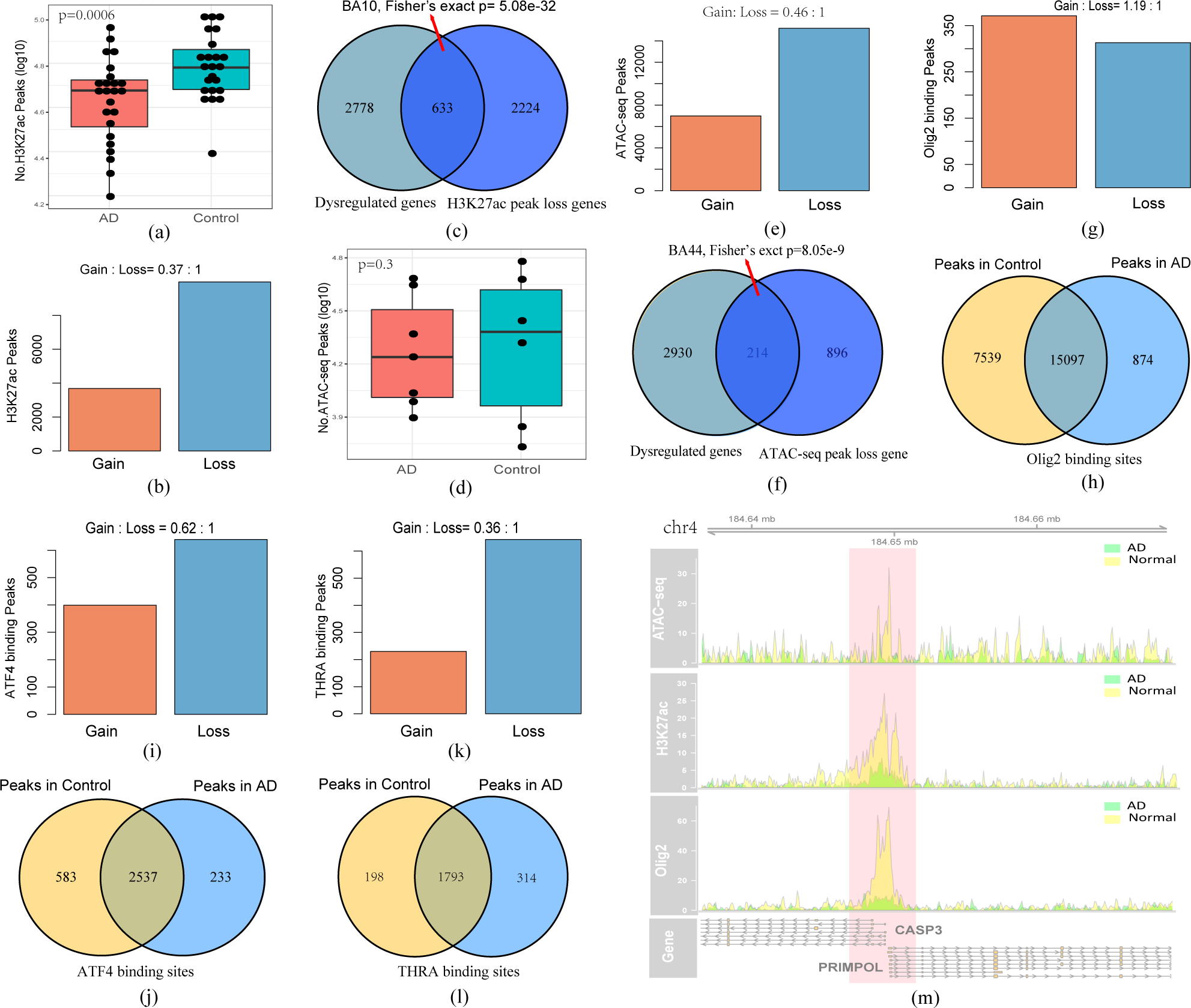
Loss of active regulation marks in AD patients. The active regulation marks are checked by differential peaking analysis and number of peaks. (a) A tendency of H3K27ac mark loss in AD patients is observed based on the peak calling analysis (*p* = 0.0006 by t-test), indicating lost activity of transcription regulation; (b) differential peaking analysis also suggests a tendency of H3K27ac mark loss in AD patients; (c) the genes with lost H3K27ac histone modification are significantly overlapped with the dysregulated genes that are identified in bi-clustering analysis; (d) ATAC-seq peaks have a weak tendency of ATAC peak loss in AD patients (*p* = 0.3); (e) differential peaking analysis to ATAC-seq data suggests more chromatin accessibility loss in AD patients; (f) the genes associated with chromatin accessibility loss are significantly overlapped with the AD dysregulated genes; (g-h) the analysis results to Olig2 binding sites, where Olig2 binding site loss is observed in AD patients; (i-l) the analysis results to ATF4 and THRA binding sites, where they have a tendency of decreased TF binding intensity in AD patient; (m) One region with the loss of open chromatin accessible region, H3K27ac mark and Olig binding site in AD patients.

Active promoters and enhancers are usually associated with open chromatin regions. We performed the assay for transposase-accessible chromatin using sequencing (ATAC-seq) to identify active regulatory regions. At a cutoff of *q*-value < 0.05, we found 207,765 open chromatin regions in 12 subjects. Both AD patients and control subjects displayed diversity in ATAC-seq peaks and there was only a weak tendency of open chromatin region loss in AD patients (*p* = 0.3) (see Figure 7(d)). We performed differential peaking analysis and found a significant tendency of open chromatin region weakening in AD patients, where the ratio of loss again gain is 1:0.46 (see Figure 7(e)), which supported a open chromatin region loss in AD patients. We then evaluated their association with dysregulated gene expression. Using the genes mentioned above, we observed a significant overlap (*p* = 8.05*e* − 9)(see Figure 7(f)), indicating that lost open chromatin accessible regions were significantly related to dysregulated gene expression.

We also performed ChIP-seq analysis to identify the binding sites of three TFs, including ATF4, OLIG2 and THRA that were predicted to be significantly associated with AD in our analysis. In Figure 7(g), we showed the result of differential peak analysis to Olig2 binding sites and found a weak binding gain in AD patients, where the ratio of binding site gain against loss is 1.19:1, which did not support regulation loss. Then, we did the peak calling analysis for each sample and observed more Olig2 binding sites missing in AD patients. As shown in Figure 7(h),7539 peaks were missed in AD patients while only 874 peaks were gained. This result suggested that Olig2 binding sites were significantly lost in AD patients. We performed the same analysis to ATF4 and THRA. Unlike Olig2, we only observed a weak ATF4 binding peak missing in AD patients.

However, a stronger tendency of binding intensity weakening was observed in differential peaking analysis for both TFs, especially for THRA (see Figure 7(i-l)). We also analysed the overlaps between lost TF binding sites and the corresponding dysregulated genes. We mapped ChIP-seq TF peaks to their nearby genes (see Figure S9). We observed significant overlaps with the lost binding sites of ATF4 (*p* = 0.01) and THRA (*p* = 0.001) (see Figure S10), which supports the contribution of lost TF binding sites.

Overall, our results indicate a tendency of active regulation loss in epigenetic regulation and TF binding sites, supporting the existence of regulation loss.

### 2.7 Ageing may contribute to the regulation loss in AD

To evaluate the impact of age on regulation loss in AD patients, we investigated if the predicted regulators displayed age-dependent expression or DNA methylation patterns. We checked predicted TFs with reported ageing-related genes [54]. But no significant enrichment was observed (*p* = 0.15), which indicated that the TFs with regulation loss were not necessarily ageing-related genes. Next, enrichment analysis was performed on the downstream target genes that were dominantly regulated by WR/MR regulators. We found that 48.8%, 52.3%, 42.3%, and 67.9% of regulation-lost regulators from four brain regions were significantly enriched with ageing-related genes (see Table S3). It seems that about half of the regulation lost TFs have regulatory roles to the ageing-related genes, which suggested a close relationship between AD and the ageing process.

We performed regulation loss study with respect to the ageing process using a set microarray expression data from human brain prefrontal cortex. This dataset included 229 AD-free participants with a broad age range from 0 to 70 years old [55]. Using the same analysis parameter, 447 regulators were predicted to be associated with ages. Like that of AD, the ageing process was accompanied with increased RLB, where Spearman’s correlation between RLB and ages is 0.779 (see Figure 8(a)), which indicated important roles of regulation loss in the ageing process. We analyzed the overlap of regulation-lost regulators from AD and ageing and found that 262 TFs were shared (see Figure 8(b)). However, this overlap was not statistically significant (*p* = 0.18). This result suggests that the regulation loss in AD patients is not the exact same regulation loss that occurs during the ageing process.

**Figure 8:**
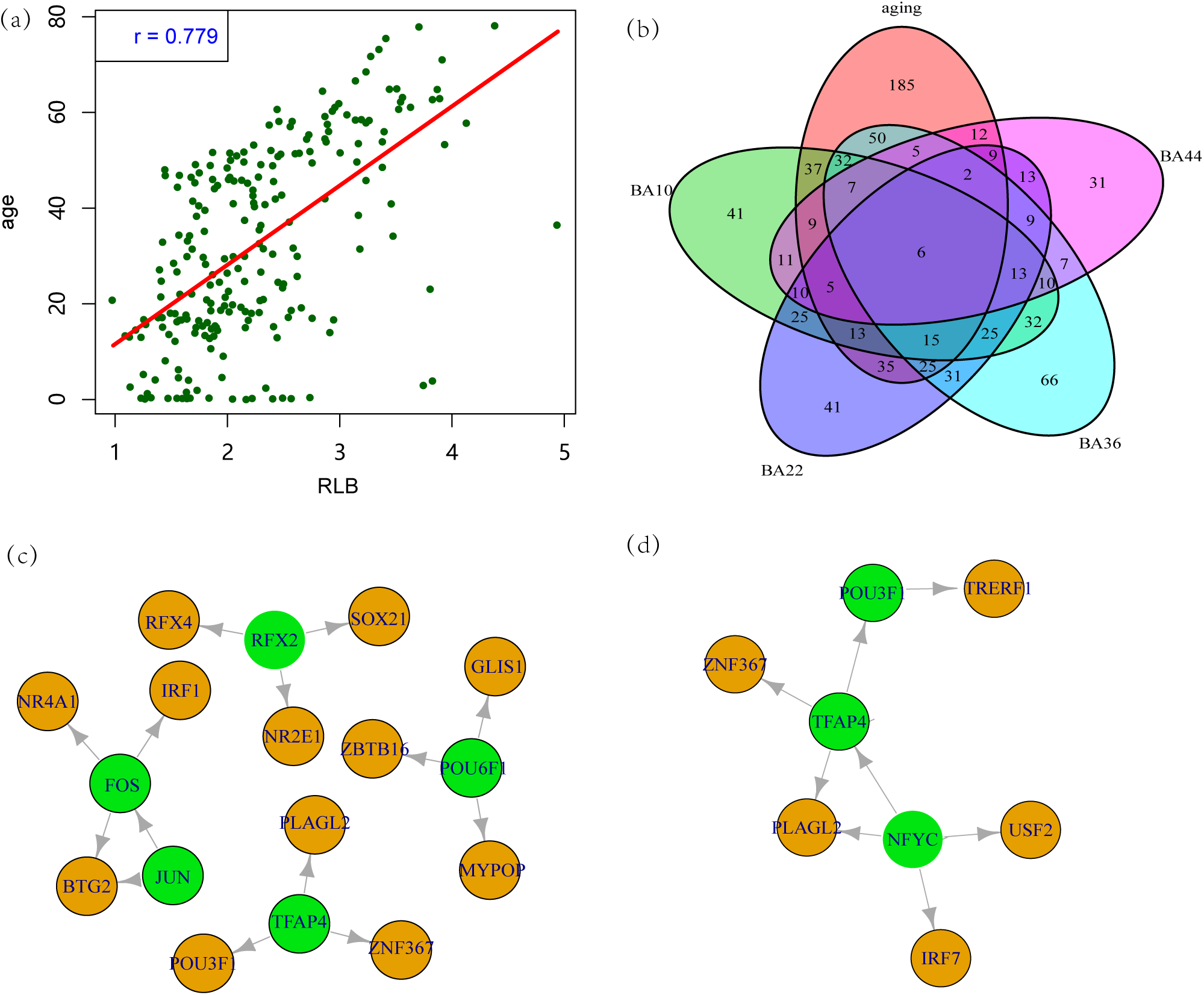
The ageing process may contribute to regulation loss in AD. (a) Regulation loss burden is well correlated with ages, indicating that regulation loss contributes to the ageing process. (b) TF overlaps among the ageing process and AD in four brain regions, which suggests the close relationship between AD and the ageing process. (c) and (d) show the regulatory network predicted from regulation loss studies, where the green nodes are regulation-lost TFs in the ageing process while brown nodes indicate TFs of AD.

We investigated the impact of ageing on AD via two manners. First, we compared the transcriptional regulatory relationships from WR and MR TFs in the ageing process to those of TFs in AD. After filtering the regulations without age correlated expression patterns, 5 regulators were identified, including three age-positive correlated TFs FOS, JUN and TFAP4, and two age-negative correlated TFs RFX2 and POU6F1. Figure 8(c) shows the relationships of these transcriptional regulators. In this network, only 16 regulators were included. However, nearly all of them were associated with AD or neuron related functions based on text-mining analysis. Another analysis was by identifying the shared WR and MR regulations by both AD and the ageing process. After filtering genes without age-dependent expression patterns, we proposed 10 regulators. In Figure 8(d), we showed the regulators and their relationship to AD regulators. By checking their expression pattern in the ageing process, we found that most of them (82%) had negative age-correlated expression profiles. It seems that age-dependent down-regulation contributes to regulation loss in AD. However, only a limited portion of AD regulation loss can be explained by this interpretation.

Overall, we found a close relationship between the ageing process and the AD process. However, it is still unclear how the ageing process affects the AD development. One hypothesis is that the regulation loss during the ageing process triggers AD genesis.

## 3 Discussion

In this work, we performed the first investigation to the diversity of AD patients in both regulation statuses and clinical manifestations. We found that accumulated degeneration of transcription regulation widely existed in AD patients and contributed to the disease development, the detrimental clinical outcomes and the ageing process. This conclusion is drawn based on results using computational modelling analysis to large-scale RNA-seq data and genome-wide studies to epigenetic marks of active regulation, where we found that (1) transcriptional regulation tends to get lost in AD patients; (2) transcription regulation loss almost indicated detrimental clinical outcomes;(3) regulation loss burden was better correlated with the clinical outcomes than the existing methods; and (4) regulation loss mainly disrupts the AD-related biological processes, especially protein degradation, neuroinflammation, mitochondrial function and synaptic function.

Recently, a growing number of epigenetic studies of AD have been described. Studies using HDAC inhibitors exert protective roles by improving dendristic spine density and by facilitating learning and memory formation [25, 56]. H3K27ac has been studied using genome-wide ChIP-seq studies. Differential peaking analysis has suggested that there is more peak loss in AD patients [24]. H4K16ac, another aging-related active transcription marks also indicated a tendency of H4K16ac mark loss [23]. Tau protein burden is reported to have a broad effect on the epigenome [19]. These findings suggests that epigenetic alterations are involved in the AD genesis and development.

Many biological processes and pathways are associated with AD, such as the presence of amyloid deposition, neurofibrillary tangles, synaptic dysfunction, neuroinflammation, neuronal cell death and oxidative stress [57]. However, the specific mechanisms underlying these AD-related dysfunctions are still not completely understood. Our present study may explain these observations as a consequence of degeneration of transcriptional regulation. We found that the participant genes of many AD-related biological processes and pathways were under dysregulation of predicted TFs. TF-mediated regulation loss can lead to their disturbance at the transcriptional level in AD patients. Among them, protein degradation, neuroinflammation and mitochondrial function are the most affected processes. They may explain the visible neuropathological features of AD patients, such as amyloid plaque and neurofibrillary tangles.

Combining these findings, it may lead to a novel causal mechanism for AD genesis and development, where brain transcriptional regulation degenerates from an organized system in normal individuals into a deficient system in the AD patients, which disrupts many AD-related biological processes. Compared to existing hypotheses, this mechanism better integrates the existing knowledge about AD under a unified framework, including the epigenetic dysregulation [17, 18, 19], disturbed gene regulatory network [9, 10, 11], broad involvement of multiple biological processes, complex clinical manifestations and AD patient diversity. AD genesis and development can be described like that epigenetic dysregulation or missing TF binding sites disrupt the brain transcription regulation and lead to abnormal cellular responses to internal or external signals; regulation loss mainly affects the AD-related biological processes, such as protein degradation, neuroinflammation mitochondrial function and synaptic function, which may contribute to the clinical outcomes of AD patients, including tau aggregations, amyloid plaques, neuroinflammation and mitochondrial dysfunction in the brain.

In the context of this work, transcriptional regulation loss indicates that TFs no longer exerts regulatory roles to their downstream genes at the right time and right genomic locations. And it is not necessary to indicate non-expression or low-expression of TF genes or the targeted genes. This is validated by the differential expression analysis between subjects with or without TF regulation, where the downstream genes of dysregulated TFs were not necessarily differentially expressed. Hence, it remains unclear as to what contributes to regulation loss in AD patients. We investigated expression changes of MR and WR regulators by differential expression analysis and differential isoform usage analysis. However, differential expression of TF genes was only observed for a few TF genes and they did not exhibit significant enrichment. As such, it is likely that regulation loss is regulated at the levels beyond the expression level of TFs. In our study of active transcription marks, e.g. H3K27ac, open chromatin accessible regions and TF binding sites, we observed a tendency of active mark loss in AD, indicating the transcription loss may result from the consequence of multiple dysregulations at different levels. Another guess is that the ageing process contributes to the regulation loss. We attempted to identify the TFs in the ageing process that also contributed to regulation loss in AD. Our analysis indicated that they were not the same process for the limited overlap of lost TFs. Furthermore, we attempted to build the connection from the lost TFs in the ageing process to the dysregulation in AD. Several TFs were predicted to take tying roles to AD. However, their contribution is too limited to account for regulation loss in AD. It is possible that the ageing process just triggers AD genesis.

Current analysis pipeline has several limitations. First, it is not surprising to observe regulation loss happened following increasing ages, which may explain the causal mechanism of the ageing process in a reasonable way. However, this also complicates our study to AD for their co-occurrence. Since age is the biggest risk factor of AD, it is important to better understand how the ageing process contributes to AD genesis. However, by now, there is still no clear clue to bridge them. In terms of the methodology of our present study, bi-clustering as a data mining method, is also an NP problem. We applied an approximating solution to identify the gene-subject combinations by gradually removing the subjects and checking improvement of Spearman’s correlations. This algorithm might reach a local optima or even a false solution. Another limitation is that we used a set of arbitrary cutoffs in our bi-clustering analysis. For example, we used |*r*| > 0.8 as the cutoff of TF-gene dominant regulation and used a minimum number of 30 genes and 50 patients as the cutoff for successful bi-clustering results. Due to the limited knowledge of TF action mechanism, the selected cutoff setting may not be in accordance with the actual scenarios. Meanwhile, the bi-clustering could generate a continuous combination of subject and genes. How to select the best gene-patient blocks is also a great challenge. We choose the combination when the number of block genes reaches the maximum values. This may not work for some TFs. In the future, we would improve the bi-clustering algorithm so that it can optimize the output based on the properties of bi-clustering output.

## 4 Data and Methods

### 4.1 Brain samples

Postmortem brain samples in the prefrontal cortex regions of 26 individuals, including 13 diagnosed with AD and 13 normal subjects were collected from the Chinese Brain Bank Center in Wuhan (CBBC, http://cbbc.scuec.edu.cn) and China Brain Bank, Zhejiang University (http://www.neuroscience.zju.edu.cn). Informed consent for autopsy has been signed for all the subjects by brain banks when the participants were in life. The clinical information of each subject was reviewed by independent neurologists with expertise in dementia and the neuropathological diagnosis was given regarding the most likely clinical diagnosis at the time of death. This study was reviewed and approved by the Ethics Committee of both brain banks and Shanghai University of Chinese Medicine for both ChIP-seq and ATAC-seq. The final sample size was limited by the available AD samples. 13 AD samples were collected and one sample was excluded for bad sample quality. We prioritized the samples with no AD or other neurological clinical manifestation as the control samples. The AD and normal samples were assigned into two groups (each size of 6): one group was used for ChIP-seq experiment of OLIG2 and ATF4 and another group was used for ChIP-seq of THRA1 and ATAC-seq. The subject information is available in Table S4.

### 4.2 Chromatin Immunoprecipitation (ChIP-seq)

The whole experiment was performed following the published protocols (see Supplementary Methods). The used the antibody includes (1) ATF4, CTS,11815S (lot#4); (2) Olig2, RnD, AF2418 (lot#UPA0718031); (3) THRA, SantaCruz, SC-56873 ((lot#J1614).

### 4.3 Assay for Transposase-Accessible Chromatin using sequencing (ATAC-seq)

ATAC-seq was performed in GENEWIZ company following the protocol introduced in [58, 59] (see Supplementary Methods).

### 4.4 Data collection and processing

The RNA-seq expression data were collected from the Mount Sinai Brain Bank (MSBB) study of Accelerating Medicines Partnership-Alzheimer’s Disease (AMP-AD) projects. The raw counts data were firstly filtered to remove the non-expressed genes by setting the maximum number of subjects with counts 0 is less than 10%. Then, the counts data were normalized by the TMM algorithm implemented in edgeR package [60]. A linear regression model was used to adjust the effects of covariates, including age, sex, postmortem interval (PMI) and RNA integrity number (RIN). The covariates were evaluated by the principal components to make sure their effects are excluded. The samples with obvious deviation were treated as outliers and removed for further analysis. The independent expression data, including ROSMAP study using microarray [9], ROSMAP study using RNA-seq [10], HBTRC study [9], Mayo’s RNAseq study for cerebellum (CBE) and temporal cortex (TCX) [52]. All of them can be retrieved from AMP-AD projects. Like the ROSMAP data, the RNA-seq counts data were collected and the similar processing pipeline was applied. For the microarray, the normalized data were collected and post-processing was performed like that of RNA-seq data.

### 4.5 Transcription factor

The TFs were selected based on the Gene Ontology (GO) annotation. The used GO biological process terms included “DNA binding activity” and “ transcription regulator activity”. 1855 genes with both GO annotation were collected. Then, they were filtered for the ones with gene expression equal to 0 in any sample of RNA-seq data. Finally, 869 TFs were used for bi-clustering analysis.

We also performed TF binding site over-representation analysis for the target genes identified by bi-clustering analysis. Here, TF binding annotation were collected from RcisTarget package [28]. 490 out of 869 TFs were selected for evaluation by filtering the ones without binding motif annotation and the ones with less than 50 target genes. Then, the TF identified by bi-clustering analysis were checked for motif enrichment. To evaluate random occurrence of enriched TF binding sites, we randomly selected *n* brain-expressed genes, where *n* was equal to the number of TF-regulated target genes, and found the chance for false recovery ratio was about 6% on overage.

### 4.6 Bi-clustering analysis

We developed a bi-clustering algorithm to study patient divergence among AD patients. The philosophy behind this algorithm was to cluster the AD patients into subsets with different TF regulation status. To measure the regulation status, we selected a set of biomarker genes to indicate the TF activity, including both average co-expression correlation and number of TF-regulated genes. To make sure of the confidence, we choose a set of strict parameters setting to define the dominant regulation, including minimum size of TF regulated patients *n* > 50, minimum number of TF-regulated genes *m* > 30 and minimum co-expression correlation *r* > 0.8. Only the result satisfying all these thresholds would be reported. As the process of bi-clustering analysis would generate a continuous number of patients and genes, the solution of bi-clustering analysis was reported under three scenarios: the solution with the maximum number of genes, the solution with the maximum number of patients and the solution with the maximum multiplied product values of patient and gene number. We evaluated them for their clinical association and found that the first one had overall better clinical consistency, which better indicated the status of regulation loss.

The full description of the algorithm is available in Supplementary Methods. The R and C++ source for bi-clustering analysis is available: https://github.com/menggf/bireg.

### 4.7 Regulation loss burden

We used regulation loss burden (RLB) to measure the degree of regulation loss in a subject. Its calculation was based on the prediction result of bi-clustering analysis. In this process, each subject was denoted by a binary vector *x*_*i*_, in which 1 indicated transcription regulation of a corresponding TF and 0 indicated no regulation. RLB is defined as

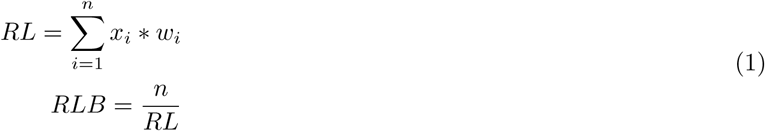

where, *n* is the number of TFs and *w*_*i*_ is a constant value that is only determined by regulation types. In this work, we tried different *w*_*i*_ values for WR TFs, ranging from 0 to 1, and evaluated them by fitting the clinical status of subjects. The evaluation results indicated the values from 0.5 to 1 acceptable in most of cases. In this work, we used the following setting:

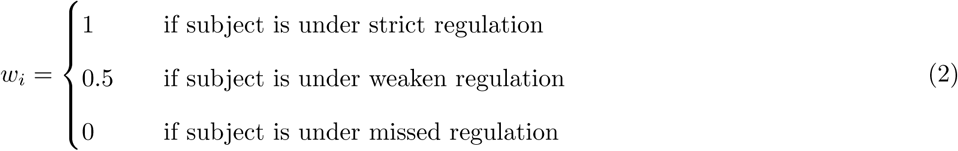

### 4.8 Peak calling

Paired-end reads of ChIP-seq and ATAC-seq DNA fragments were generated by Illumina Novaseq 6000 platform. On average, 20 million reads were available for each sample. The raw fastq files were firstly checked by *fastqc* and found overall good quality of the sequencing data. We used NGmerge to remove the adaptors [61]. Then, reads were aligned to human genome hg38 using bowtie2 [62]. The output sam files were filtered for reads with PCR duplicates, unmapped tags, non-uniquely mapped tags or with mismatches greater than 2. SAM files were then transformed into sorted and indexed bam files by samtools [63]. We applied macs2 [64] to identify the peaks under a cutoff of *q* < 0.05. For ATAC-seq data, we used a set of option “–broad –nomodel –shift 37 –extsize 73 –keep-dup all”.

For both AD and control group, we observed that the number of reported peaks are quite diverse among subjects. To identify the peaks with more confidence, we did cross-validation to select the peaks observed in more than one sample using “bedtools”. Each “*.bed” file was checked with other peak files using “bedtools intersect -wa -f -r 0.6”.

### 4.9 Differential peak analysis

Besides of peak calling analysis for each sample, we also did differential peak analysis to study the overall histone modification/open chromatin accessibility/TF binding intensity in AD and control samples. In this process, we applied an analysis pipeline proposed by S.J. Marzi et.al [24]. In peak calling steps, bam files of both AD and normal subjects were merged to maximize the power of peak calling. Under a parameter setting mentioned above, MACS2 [64] was used. Considering the fact that the histone modification marks usually spread broad regions, we set “–broad” option for the ChIP-seq data of H3K27ac, H4K16ac and ATAC-seq data. We validated and filtered the identified peaks by checking the peak overlap using the ones identified in the NIH RoadMap Epigenomics Consortium in all brain regions, including angular gyrus, anterior caudate, cingulate gyrus, BI.middle hippocampus, inferior temporal lobe, middle frontal lobe and substantia nigra. Next, we estimated the read abundance of peaks using the tools in `Rsubread` package. After filtering the peaks with read count of 0 or total reads number is less than 100, we did differential peak analysis to find the genomic regions with regulation loss and gain. In this step, we used edgeR [60] to identify the peaks with altered TF binding affinity or open chromatin accessibility status. Different from S.J. Marzi et.al’s analysis pipeline, we consider the facts that the peaks loss and gain are not equal in AD samples and one-step differential peak analysis may cause under-estimation to the regulation loss. A better solution is to normalize the count data using only peaks without regulation loss or gain. Therefore, we applied a multiple round analysis. In each round, we perform differential peak analysis and removing the differential peaks. This process was repeated until no new differential peak was reported anymore. The library size and dispersion information of count data were updated using the count matrix reported in last round of differential peak analysis. Considering that the selected histone modification markers, open chromatin accessibility and TF binding sites were all associated with active regulation, we reported the peaks with increased read abundance as regulation gain and the peaks with decreased abundance as the regulation loss at a cutoff of FDR < 0.05. Using `ChIPseeker` package [65], we annotated the differential peaks for their genome location and nearby genes in a arrange from −3000 bp to 1000 bp round transcription start sites.

The perl and R source code for differential peak analysis is available:https://github.com/menggf/bireg/tree/master/code.

### 4.10 Functional annotation

Our evaluation suggested that the regulators took dominant regulatory roles to their downstream genes. The functional involvement of regulators was inferred by annotation to the downstream regulated genes. The R package `clusterProfiler` was implemented for enrichment analysis using Gene Ontology biological process terms. Under a cutoff of Benjamini *p* < 0.05, the enriched terms were selected and calculated for their frequency among all the regulators. After manually filtering the terms with overlapped functional annotation, terms with both significant enrichment and high frequency were identified to represent the functional involvement of regulators. The used terms were clustering using the hierarchical clustering method based on the *p*-values reported by enrichment analysis. To smooth the heatmap visualization, the *p*-values were transformed with −*log*10. If the −*log*10(*p*) value is greater than 6, they were set to 6. We also did the functional annotation for each regulator by text-mining tools, including Ingenuity Pathway analysis (IPA), GeneCards (https://www.genecards.org/) and PumMed, to evaluate the functional consistency among prediction from different sources.

### 4.11 Enrichment analysis

Enrichment analysis was performed to evaluate the statistical significance of feature overlaps, such as genes associated with the same biological process. For *k* input genes, the number of genes with certain feature is *x*. Among *n* whole genomic genes, the number of genes with such feature is *p*. Fisher’s exact test can evaluate if the observed *x* genes resulted from random occurrences. We used the R codes to calculate statistical significance:

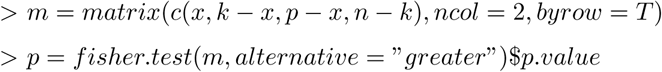

### 4.12 Co-expression network analysis

We used WGCNA, as the implementation of co-expression network analysis, to predict gene modules of expression data by following the protocol introduced by the official document in https://labs.genetics.ucla.edu/horvath/htdocs/CoexpressionNetwork/Rpackages/WGCNA/Tutorials/index.html. Considering the fact that WGCNA has some parameter setting while no golden standard to optimize them, we test different setting one by one and chose the setting combination where best clinical association were observed as the final parameter setting. The summarized profiles or eigengenes of modules were investigated for their clinical association by Spearman’s correlation. Except for the gray module, modules with the best clinical correlation were selected for evaluation.

## Supporting information

Supplementary Results

## 5 Data Available

The ATAC-seq and ChIP-seq data were publicly available in the Gene Expression Omnibus (GEO) database with the ID of GSE129041. Its reviewer token is: epkfqwoyzfsdxed.

## 6 Acknowledgements

We thank GSK R&D China in Shanghai (closed in 2018.04) for supporting this research work in Alzheimer’s disease. We are grateful to the China Brain Bank of Zhejiang University School of Medicine and Chinese Brain Bank Center in Wuhan for providing human brain material. The results published here are in part based on data obtained from the AMP-AD Knowledge Portal (doi:10.7303/syn2580853). We appreciate their generous contribution to the studies of Alzhiemer’s disease. We thank Dr. Lichun Jiang, DR. Ruping Sun for reading this manuscript and giving us many useful suggestions. We also thank Dr. Guhan Nagappan, Dr. Xiaoming Guan, DR. Bai Lu for valuable comments.

